# Sequence alignment of the primate lineage reveals evolutionary divergence and conserved secondary structural motifs in noncoding RNAs

**DOI:** 10.64898/2026.05.18.725462

**Authors:** Anish Beeram, Zion R. Perry, Anna Marie Pyle

## Abstract

Long noncoding RNAs (lncRNAs) constitute most of the human transcriptome and perform essential roles in chromatin organization and transcriptional regulation. Because lncRNA genes are not constrained by protein-coding ability, they tend to exhibit more rapid evolutionary divergence. Their poor nucleotide sequence conservation among mammals often led to the assumption that lncRNAs lack conserved structures. However, emerging evidence indicates that many noncoding RNAs adopt secondary and tertiary folds critical for protein recruitment, chromatin binding, and regulation of gene expression. Nevertheless, there are few experimental secondary structures for lncRNAs, hindering mechanistic insight into lncRNA structure-function relationships. Even without available structural data, covariation, in which two nucleotides co-evolve, can provide evidence for conserved structures. This requires sequence alignments with sufficient divergence to detect covariation but enough similarity to maintain alignment quality. Here we report the development of a novel computational pipeline to mine 190 unannotated primate genomes to generate high-quality multiple sequence alignments of noncoding RNAs. This pipeline performs sequence searching, locus extraction, cross-species alignment, and downstream analyses, including assessment of covariation and primary sequence conservation. Ultimately, we demonstrate that because many noncoding elements, such as lncRNAs evolve at a more rapid rate than protein-coding genes, phylogenetic analyses constrained within a narrower evolutionary span can be used to identify conservation of primary sequence and secondary structure. By focusing our alignments on the primate lineage, our method overcomes the limitations of broad phylogenetic analyses, enabling high-resolution detection of subtle conservation patterns and conserved secondary structural motifs of long noncoding RNAs.

## INTRODUCTION

Long noncoding RNAs (lncRNAs) are RNA transcripts longer than approximately 500 nucleotides that generally do not encode proteins (Mattick et al. 2023). While protein-coding genes make up only a small fraction of the human genome, lncRNA genes are remarkably numerous—current GENCODE annotations list roughly 19,000 protein-coding genes but over 35,000 lncRNA genes in humans (GENCODE 2025).

Importantly, lncRNAs have emerged as crucial regulators of gene expression and cellular physiology. The XIST lncRNA (∼17 kb) is the driving regulator of X-chromosome inactivation, recruiting chromatin-modifying complexes to silence gene expression (Brown et al. 1992). Other well-studied examples include HOTAIR, which scaffolds PRC2 and LSD1 to modulate chromatin state (Rinn et al. 2007; Somarowthu et al. 2015), and MALAT1, which has yielded pro-metastatic phenotypes in lung adenocarcinoma models due to transcriptional upregulation of metastasis-associated genes including GPC6, LPHN2, and CDCP1 (Gutschner et al. 2013).

LncRNAs are often described as exhibiting rapid evolutionary divergence and poor primary sequence conservation (Kutter et al. 2012; Szcześniak et al. 2021). Unlike protein-coding genes that are constrained by the genetic code, lncRNA sequences accumulate mutations more frequently and commonly lack apparent one-to-one orthologs across distant species (Quinn et al. 2016). This limited conservation historically led to the belief that lncRNAs might lack meaningful structure or function. As such, early covariance analyses—which evaluate whether coordinated substitutions preserve base-pairing and secondary structure—found little evidence of conserved base-pairing in well-known lncRNAs such as XIST and HOTAIR (Rivas et al. 2017; Tavares et al. 2019).

Nevertheless, structural studies have revealed that several lncRNAs adopt complex conformations essential for function. For example, the RepA region of XIST folds into three independently folding domains with conserved tertiary structure (Liu et al. 2017). In lincRNA-p21, inverted repeat Alu elements form a conserved double loop structure necessary for nuclear localization and retention (Chillón and Pyle 2016). The 3’ stem-loop of MALAT1 has also been observed to form a triple-helix structure that inhibits nuclear degradation of the RNA (Brown et al. 2012; Brown et al. 2014).

Despite these advances, understanding of lncRNA structure–function relationships remains limited. Only a small number of lncRNA secondary structures have been experimentally resolved, leaving most human lncRNAs structurally ambiguous (Novikova et al. 2012). This gap stems from technical challenges: lncRNAs are typically large and contain both well-folded and highly disordered regions, making crystallography or NMR studies challenging; they are expressed at low abundance, have alternative isoforms, and are often bound to ribonucleoproteins, complicating purification of lncRNA in their native-folded states (Wu et al. 2021; Ouyang et al. 2022; Chen and Kim 2024). As a result, there are only a few high-quality secondary structures of lncRNAs in cells (Sherpa et al. 2018; Uroda et al. 2019; Monroy-Eklund et al. 2023; Falese et al. 2025; Oh et al. 2025). Compounding this limitation, few studies have examined sequence divergence of noncoding RNAs within relevant evolutionary time scales and whether there is support for conserved structural motifs (Sigova et al. 2013; Hezroni et al. 2015).

Bridging this structure-function gap is crucial. Here we provide an *in silico* approach for the determination of conserved primary sequence and secondary structural motifs in ncRNAs. Our novel computational approach mines and analyzes unannotated, un-scaffolded raw sequencing data from 190 primate genomes to generate high-coverage sequence alignments of ncRNAs (Kuderna et al. 2023). Because of the rapid sequence divergence of long noncoding RNAs, we hypothesized that narrowing the scope of our pipeline to the primate lineage would enable the detection of conservation and covariation in lncRNAs. This will allow the identification of secondary structural motifs with strong evolutionary support for functional significance. The purpose of this study is to benchmark and validate this novel tool for determining conservation in both primary sequence and secondary structure using candidate noncoding RNAs for which experimental structural data is readily available. We chose 2 untranslated regions (UTRs) and 1 lncRNA for which *in cellulo* secondary structural data exists. We found highly conserved regions of the primary sequence in alignments of the candidate lncRNA and both control UTRs. We evaluated conservation of ncRNAs across the primate clades to analyze their patterns of evolutionary divergence, and subsequently demonstrated that these alignments have the power to detect covariant base-pairing, identifying conserved, functionally relevant secondary structural motifs in selected lncRNA helices. These findings suggest that our novel bioinformatics pipeline can identify putative ncRNA loci in unannotated primate genome assemblies, rendering this raw sequencing data interpretable for evolutionary analyses of RNA. For any RNA or genomic loci of interest, this pipeline ultimately allows for the mapping of conservation patterns and the detection of significant covariation within the primate lineage.

## RESULTS AND DISCUSSION

### Development of pipeline for ncRNA alignment and analysis

We have developed a novel computational pipeline that mines 190 unannotated primate genomes from the European Nucleotide Archive to generate high-coverage multiple sequence alignments (MSAs) of ncRNAs (Kuderna et al. 2023). Because these genomes are unannotated and incompletely scaffolded, often just a sequence of contigs, query-to-target pairwise alignment of nucleotides was conducted using Exonerate, a command-line sequence alignment tool. This generated hundreds of candidate loci in each target genome. We selected the Exonerate alignment with the highest alignment score—indicating the highest quality pairwise alignment—before transcribing the DNA nucleotide sequence into the corresponding RNA sequence.

After collating all selected RNA sequences, an MSA was generated using the command-line tool Multiple Alignment using Fast Fourier Transform (MAFFT) with sequences aligned against the query human sequence (**Fig. 1**). The MSA could be further refined through clustering by sequence similarity with the tool CD-HIT-EST. The generated MSAs served as a basis to assess conservation and covariation using statistical tools such as RNA Structural Covariation Above Phylogenetic Expectation (R-scape) to identify candidate functional structures (Rivas et al. 2017). At present, we are the first group to mine these primate genomes for structural and evolutionary analysis of ncRNAs.

**Figure 1.**
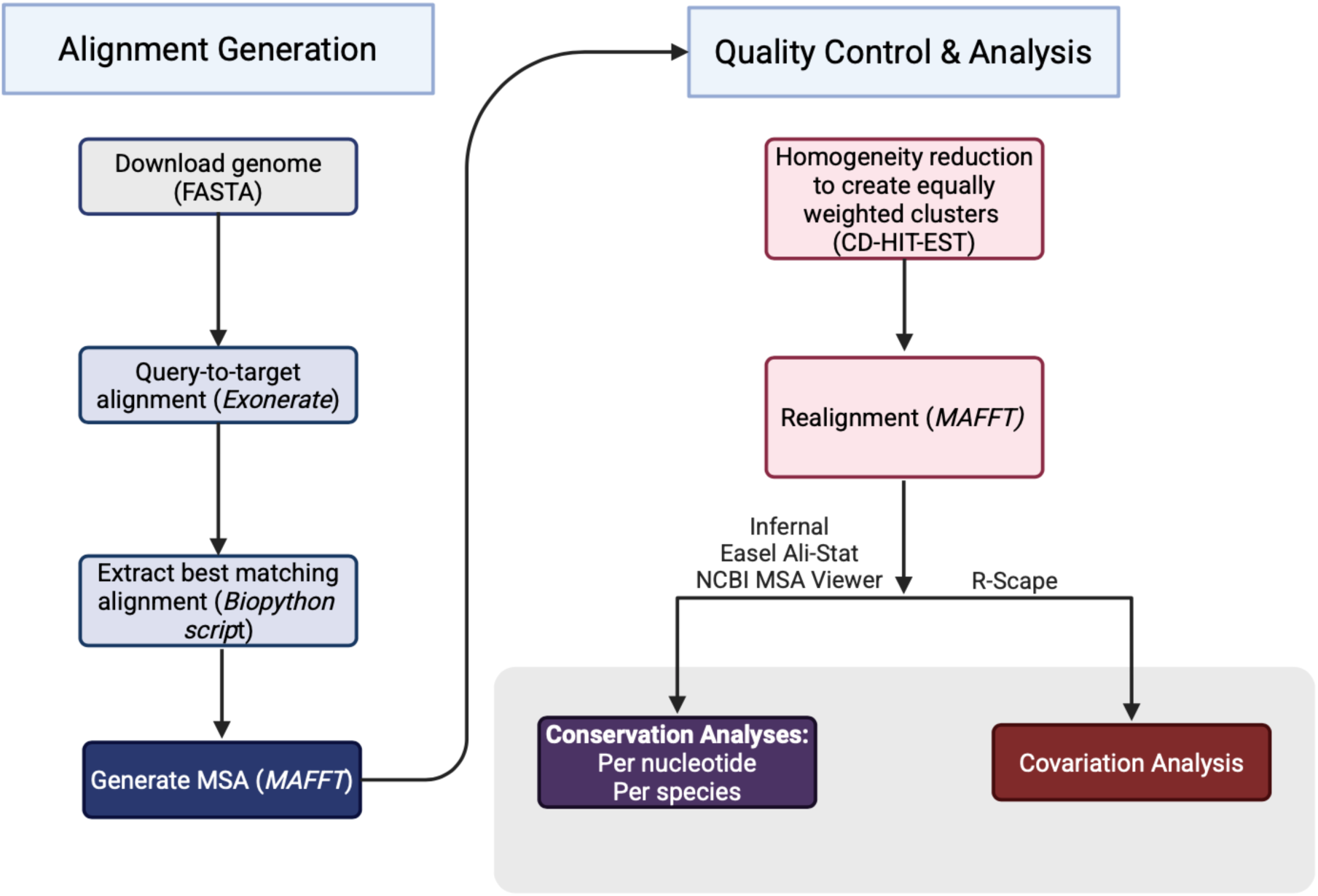
Schematic diagram to identify noncoding RNA in unannotated primate genome assemblies. Multiple sequence alignments (MSA) are generated by downloading a directory of 190 primate genomes. Query-to-target alignment is conducted with Exonerate before selecting best matching candidate RNA via custom Biopython script. MSAs are resolved with MAFFT. Homogeneity reduction is conducted to parse alignments with CD-HIT-EST before realigning with MAFFT. Conservation is determined per-nucleotide with Infernal and per-species with NCBI MSA Viewer. Covariation is analyzed with R-scape (RAFSp). Figure created in BioRender.

### Benchmarking the pipeline on conserved UTRs

The pipeline was first evaluated on 190 available primate genomes from the European Nucleotide Archive using selected ncRNAs. We chose the 5’ UTR of human TP53 (hTP53) and the 5’ UTR of human Ferritin (hFerritin), both of which have well-established structural and functional significance. The 5’ UTR of hTP53 spans 152 nts and is characterized by two highly-conserved hairpin motifs which comprise the IRES (Kim et al. 2025). The 5’ UTR of hFerritin spans 209 nts and contains the well-documented Iron Responsive Element (IRE) (Wang et al. 1990; Kikinis et al. 1995).

The 5’ UTR of P53 has been demonstrated to scaffold protein-binders including the E3 ubiquitin ligase MDM2, RPL26, and Nucleolin after being stabilized by endogenous divalent free Mg^2+^ cations (Chen and Kastan 2010; Chinnam et al. 2022; Kim et al. 2025). The 5’ IRE of Ferritin adopts a single stem-loop motif configuration which is necessary for binding of Iron Regulatory Proteins under differential iron concentrations (Harrell et al. 1991; Wang et al. 2021). These ncRNAs are ideal starting candidates, as they are well-documented, functionally important RNAs with high confidence structural data, which can be used to benchmark our pipeline on the 190-primate genome dataset.

After assembling the 190-genome MSA for each RNA candidate, per-nucleotide conservation scores were calculated using the third-party tool Infernal for Easel (esl-alistat), which quantifies on a scale from 0, indicating no conservation, to 2, which reflects perfect nucleotide conservation (Eddy 2023). We also calculated the percent non-gapped, which quantifies how much of any given alignment column consists of actual nucleotides, rather than gapped sequences which are characteristic of insertions or deletions. These two parameters indicate our confidence in locus identification by measuring how high-coverage the alignment is and level of conservation across primate species. Analysis of candidate RNA conservation and coverage revealed that alignments frequently contained multiple high-gapped regions, almost always due to insertions or low coverage alignment reads found in a small percentage of primate species—but notably, not found in the primary sequence of the human ncRNA. This finding indicated that the MSAs contained non-reference, sequence-specific insertion variants, which made total alignment per-column conservation data noisy. To determine primary sequence conservation for only nucleotide positions present in the human consensus sequence, we excluded per-column conservation and coverage of any alignment column not containing a nucleotide in the human ncRNA. In doing so, our data reflects the extent and quality of primary sequence conservation of human noncoding RNA across 190 primate species.

The primary sequences of the selected 5’ UTRs were found to have high conservation and high coverage (**Fig. 2A, B**). Median conservation and coverage of the TP53 primary sequence were 1.45/2 and 95.29%, as well as 1.87/2 and 99.47% for the 5’ UTR of Ferritin. The TP53 5’ UTR alignment is slightly less conserved throughout its transcript than the 5’ UTR of Ferritin, and notably, the coverage drops off near the end of the TP53 transcript where the final 18 nts are missing from ∼40% of species. These results indicate that both UTR alignments were high-quality, and that the primary sequences of both Ferritin and TP53 are well-conserved across primate species. Although there are few large-scale evolutionary analyses of UTR conservation across mammals, the findings with these two genes might be expected due to the high functional relevance of UTRs in translation initiation. Given their functional importance for protein-coding genes, there may be selection pressure on the primary sequence of 5’ UTRs (Churbanov et al. 2005; Byeon et al. 2021; Chaldebas et al. 2026). The Ferritin and TP53 alignments serve as positive controls for our study and confirm that specific noncoding RNA sequences could be successfully identified from 190 unannotated primate genomes.

**Figure 2.**
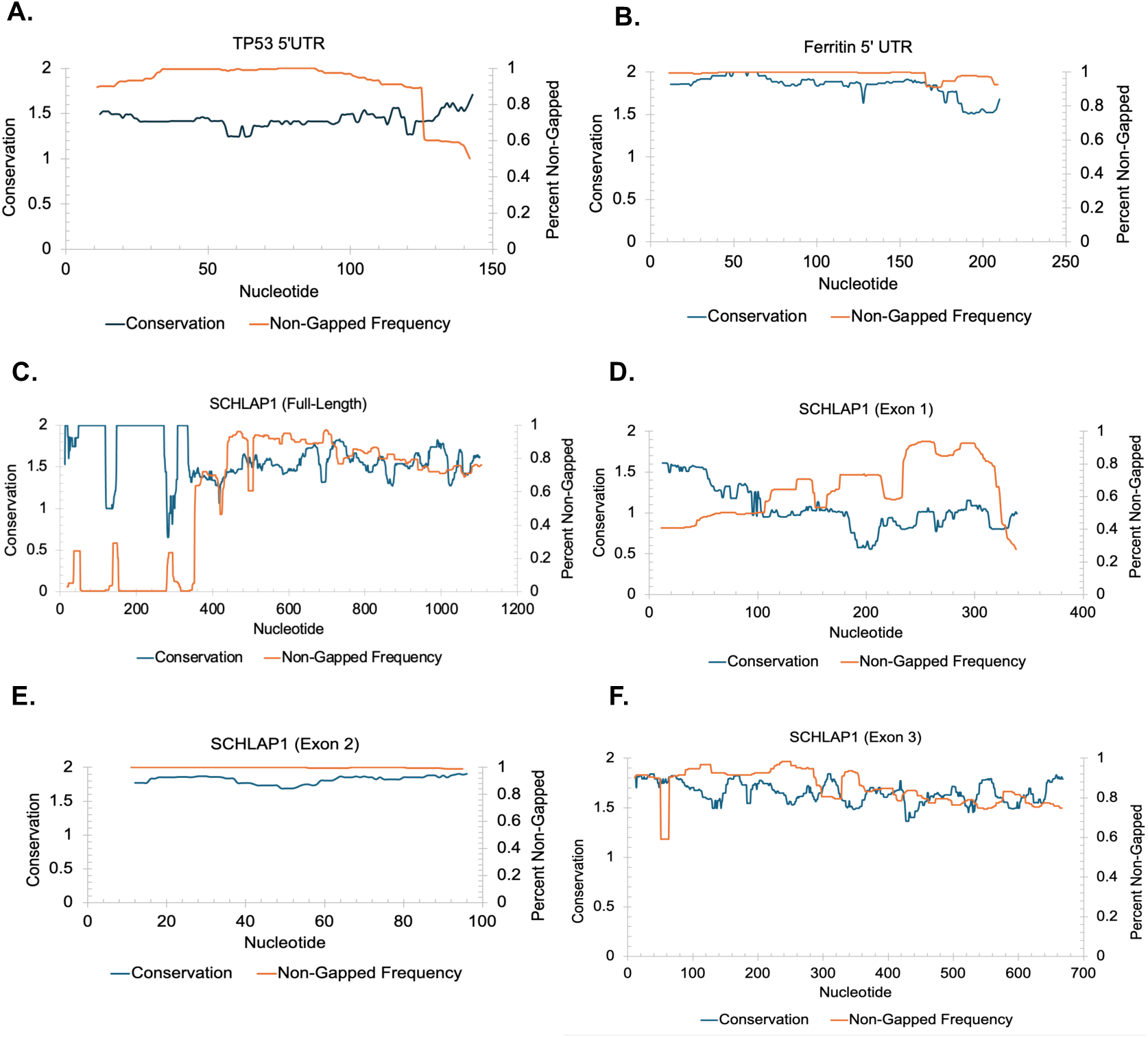
Per-nucleotide conservation for control UTRs and lncRNA SCHLAP1 validates pipeline design. Per-nucleotide conservation and non-gapped frequency plotted against nucleotide position in the human consensus sequence using a 21-nt median sliding window. (A) TP53 5’ UTR. (B) Ferritin 5’ UTR containing the Iron Responsive Element. (C) Full-length SCHLAP1 containing all three exons. (D-F) Per-exon alignments of SCHLAP1.

### Extending the pipeline to analyze conservation of a lncRNA

The next candidate ncRNA we selected was full-length human SChLAP1 (hSChLAP1, transcript variant 3) which contains 3 exons across 1.1 kb. hSChLAP1, which functions as a negative regulator of the SWI/SNF chromatin-remodeling complex, has been demonstrated to induce pro-metastatic phenotypes in prostate cancer, and recently had the secondary structure resolved *in vitro* and *in cellulo* (Prensner et al. 2013; Falese et al. 2025; Oh et al. 2025). After assembling the 190-genome MSA for hSChLAP1, including for each-exon of hSChLAP1, we conducted analyses of primary sequence conservation and coverage of the full-length and per-exon alignments.

The primary sequence of full-length SChLAP1 was found to be much more variable compared to the UTRs, as expected, but there is strong evidence for conservation within portions of the transcript (**Fig. 2C**). The first 340 nucleotides of the RNA were not identified in the full-length alignment of SChLAP1, as the median percent-coverage across this nucleotide range was 0.523%. Since the coverage is so low, the conservation score is not meaningful, since fewer than 10/190 primate species had a nucleotide present in the first 340 alignment columns corresponding to a nucleotide in the human sequence. However, the last 760 nucleotides of the alignment had starkly improved coverage and conservation, with a median percent coverage of 80.62% and conservation score of 1.54/2.

These findings led us to hypothesize that such low coverage across the first 340 nucleotides of the human sequence may be due to large introns separating exons of SChLAP1, which may hinder the accuracy of our pipeline when conducting query-to-target alignment in unannotated genomes. The SChLAP1 variant we used contains 3 exons from 1-338nt, 339-433nt, and 434-1100nt. Based on the full-length alignment (**Fig. 2C**), the pipeline did not identify the first exon in most species, had poor recognition of exon 2 (68.59% median coverage, 1.36/2 median conservation), but successfully identified the third exon (83.77% median coverage, 1.56/2 median conservation).

To determine whether exon boundaries may impede the accuracy of our alignment tools, we conducted per-exon alignments of SChLAP1 (**Fig. 2D-F**). The primary sequence of exon 1 was found to have a median percent coverage of 61.78% and median conservation of 1.05/2. While this data is noisier than our UTR alignments, these results suggest that we extracted exon 1 across 190 primates. Exon 2 has significantly improved alignment quality, with median percent-coverage of 100% and median conservation of 1.83/2. Lastly, the alignment of Exon 3 alone, like that of full-length SChLAP1, seems to have accurately captured the third exon, with median coverage of 84.15% and median conservation of 1.59/2, which closely corresponds to the quality of the full-length alignment. Taken together, the per-exon alignments of SChLAP1 produced significantly better alignments than that of full-length SChLAP1, suggesting that query-to-target alignment performs better without searching across potentially large introns.

### Comparison of evolutionary divergence of noncoding RNAs across primates

Additionally, we calculated the rate and extent to which these noncoding RNA sequences diverge over evolutionary time within primates. This allowed us to evaluate differential conservation patterns among different ncRNAs and their association with phylogenetic distance. To accomplish this, the pairwise conservation score of each species’ alignment against the human sequence was calculated using NCBI MSA Viewer to determine percent coverage and percent identity of the entire alignment, including all alignment columns and species-specific insertions. Rather than analyzing individual nucleotide position in the alignment as done in Figure 2, identity and coverage scores were assigned to entire alignments based on their total percent-similarity and percent-coverage compared against the human sequence. An effective identity score was then calculated by multiplying the percent coverage by percent identity to adequately reflect each pairwise alignment. Subsequently, the open-source database TimeTree was used to obtain the divergence time of each species relative to human (Kumar et al. 2022). The 190 species were then clustered into 5 bins based on their divergence time from human, as measured in millions of years ago (MYA): 19, 29, 43, 69, and 74 MYA. This allowed us to map each species’ effective identity against divergence time to assess the rate at which RNAs diverge across the 190 primates (**Fig. 3**).

**Figure 3.**
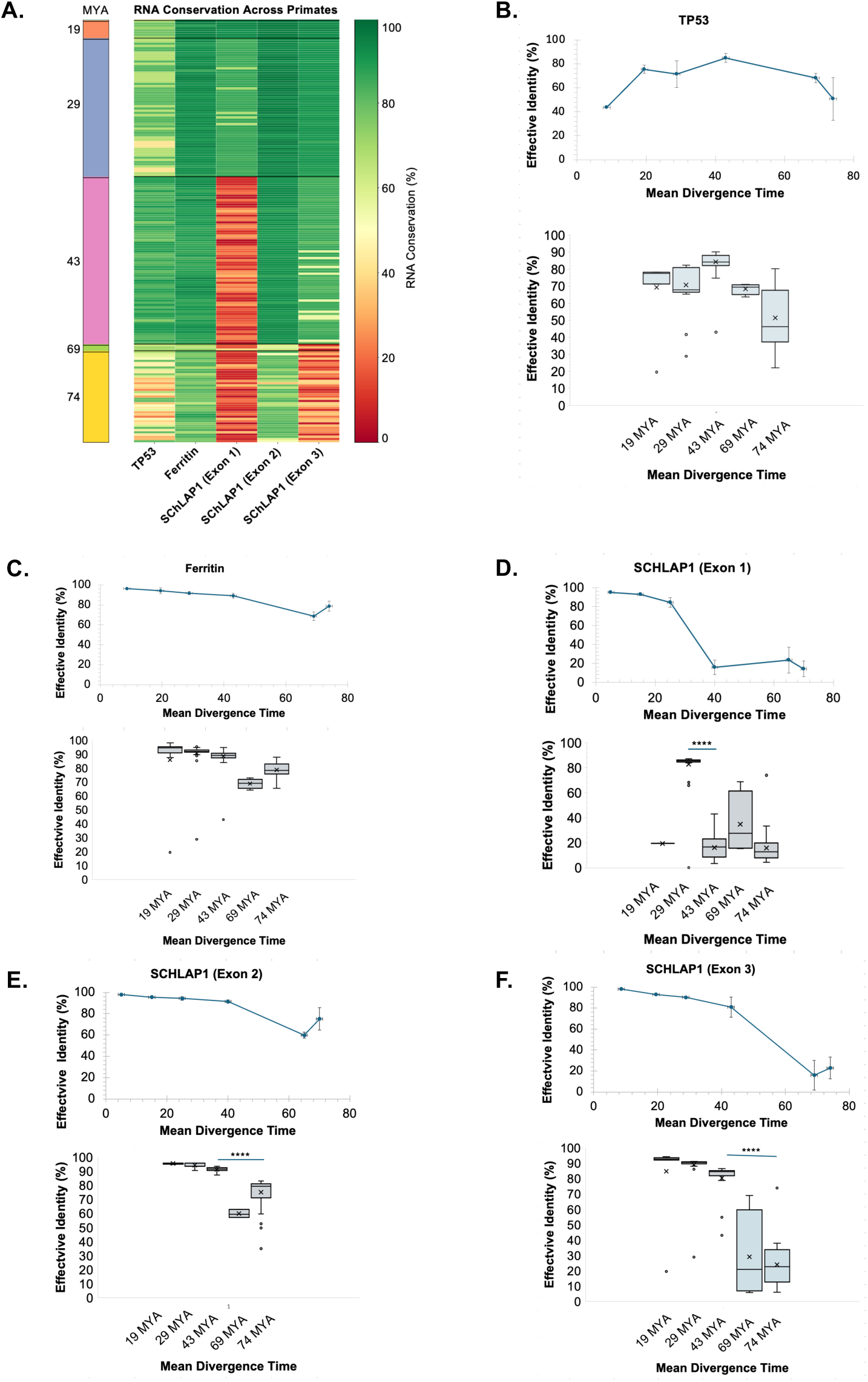
Evolutionary divergence of noncoding RNA across primate genomes by per-species pairwise conservation. (A) Heatmap depicting effective identity of each RNA per-species, sorted in descending order of divergence. (B-F) Average identity-divergence relationship and box-and-whisker plots for TP53 5’UTR, Ferritin 5’ UTR, and per-exon alignments of SCHLAP1.

The 5’ UTRs of Ferritin and TP53, which are thought to be widely conserved, indeed demonstrated little evolutionary divergence (**Fig. 3A-C**). For example, the divergence patterns for TP53 5’UTR indicated that the RNA was conserved across all five divergence bins, without a statistically significant drop in per-species conservation even as the species increased in evolutionary distance from human. Additionally, just as the MSA for TP53 yielded more variable conservation (**Fig. 2A**), the same was seen in divergence bins of 29 and 74 MYA, where TP53 effective identity scores had a larger spread with more notable outliers when compared with that of Ferritin (**Fig. 3B, C**). Even in the case of Ferritin, the lowest average effective identity score was still 68.7% similarity to the human sequence. Taken together, these UTRs serve as positive controls for RNA which are well-conserved across evolutionary time.

Notably, for the alignments of SChLAP1 exons 1 and 3, species’ conservation decreased significantly with increasing evolutionary time, particularly around 43 MYA for the alignment of Exon 1 and 69 MYA for that of Exon 3 (**Fig. 3D, F**). SChLAP1 Exon 2 seemed to be largely conserved across all 190 primates included in the MSA (**Fig. 3E**). Because the lncRNA alignments all exhibited a statistically significant decrease (**** = p ≤ 0.0001 by unpaired, two-tailed Student’s t-test) in average effective percent identity as species became increasingly more phylogenetically divergent from humans, this data suggests that it is possible to capture rapid evolution rates of long noncoding RNAs within the narrow evolutionary span of 74 MYA. Since lncRNAs are not constrained by the genetic code as are protein-coding genes, and evolve much more rapidly as a result, phylogenetic analyses must be conducted within a small enough time frame to capture and visualize the accelerated sequence changes of these RNAs. This data suggests that while mammalian alignments may be too broad of an evolutionary span, rapidly evolving genes can be analyzed at higher-resolution within the primate lineage (Johnsson et al. 2014).

### Detection of covariation in primate alignments supports functional secondary structures in SChLAP1

To discover whether there were any functionally relevant structures in the candidate lncRNA SChLAP1, we evaluated covariance using R-scape (v0.2.1, RAFSp). First, it was necessary for us to create more balanced alignments, with equally weighted sequence similarity clusters. Because the MSAs often contained a much greater number of primate genomes from 29 and 43 MYA, potential compensatory single-nucleotide polymorphisms which maintain base pairing may not be detectable by R-scape if alignments are too homogenous. To perform a homogeneity reduction, we used the command-line tool CD-HIT-EST at a similarity index threshold of 95% to create equally weighted clusters and build representative alignments.

Subsequently, we provided a secondary structure of SChLAP1 along with the alignment to enable detection of base-pairing interactions with experimental support. This secondary structure allows for orthogonal verification of the covariant base pairs found using R-scape, as R-scape reports both covariant base pairs that are supported by the MSA *and* those covariant base pairs found in the MSA which are also found in the provided secondary structure. The ratio of covariant pairs that are supported by the MSA and secondary structure (“true positive covariants”) divided by the total number of covariant pairs identified from the MSA (“covariant BPs”) is referred to as the *precision* of the covariance model (**Fig. 4D**). Therefore, we obtained *in cellulo* SHAPE-MaP reactivities for secondary structure probing of SChLAP1 (Falese et al. 2025) and predicted the secondary structure using SuperFold. After aligning the secondary structure to the MSA, we performed covariance analysis on the full-length and per-exon alignments of SChLAP1 (**Fig. 4**).

**Figure 4.**
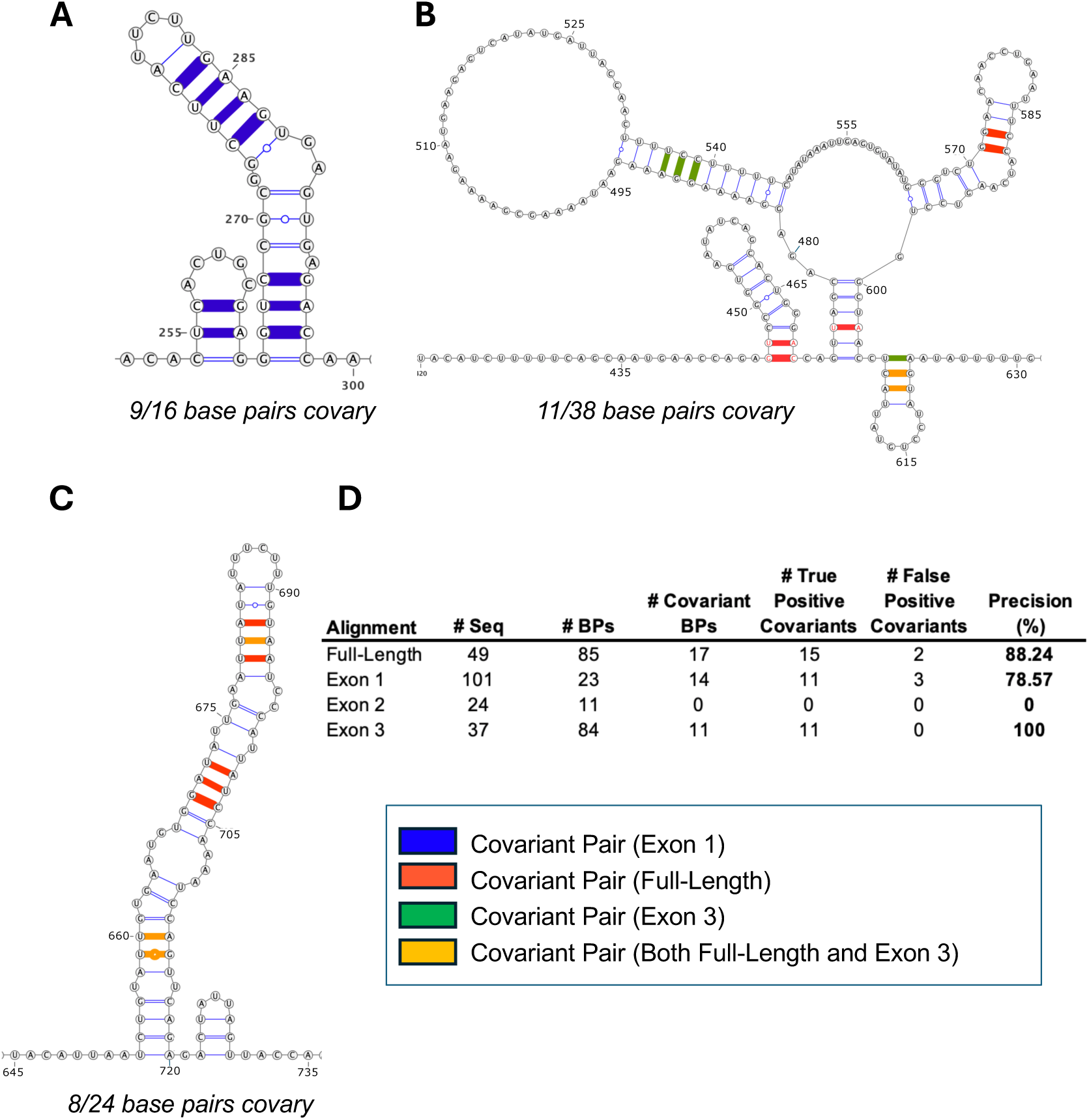
Primate alignments provide evidence for conserved secondary structural motifs in SCHLAP1. Covariation in base-paired helices of SCHLAP1. (A-C) Analysis of covariant base pairs in full-length and per-exon alignments of SCHLAP1 using R-scape (0.2.1, RAFSp), validated against *in cellulo* SHAPE-MaP probing of SCHLAP1 secondary structure (Falese et al. 2025). Covarying residues mapped onto helices in reference structure. Covariant pairs are colored according to their corresponding alignment. (D) Comparison of covariant base pairs across alignments. Precision is calculated by R-scape as a percentage of covariant base pairs identified in the MSA which are supported by the provided secondary structure.

We identified several helices in SChLAP1 (**Fig. 4A-C**) with statistically significant covariant base pairing—where the covariance was identified in the MSAs *and* subsequently cross-verified as being base-paired in the *in cellulo* SHAPE-MaP secondary structure. We highlighted the covariant base pairs on notable helical stems, indicating those identified in the per-exon and/or full-length SChLAP1 alignments. We identified 2 adjacent stem loops in exon 1 from nucleotides 254-298 for which our alignments indicated 9 covariant pairs out of 16 total base pairs (**Fig. 4A**). In exon 3, we also identified 2 regions with significant covariance: the first from nucleotides 446-622, for which 11/38 base pairs covary across 2 stem loops and a multi-helical junction (**Fig. 4B**); the second from nucleotides 653-720, for which 8/24 base pairs covary in one, long stem-loop (**Fig. 4C**). Notably, no covariance was detected in exon 2, likely due to the highly conserved nature of the alignment, which was not conducive for identifying compensatory mutations that preserved base pairing.

However, the full-length alignment of SChLAP1, despite having lower conservation scores (**Fig. 2C**) revealed 17 covariant base pairs, 15 of which were substantiated by the provided secondary structure (88.24% precision; **Fig. 4D**). The alignment of exon 1 yielded covariant base pairs with 78.57% precision and the alignment of exon 3 had 100% precision, since all 11 covariant base pairs were base-paired nucleotides in the secondary structure (**Fig. 4D**). Taken together, this data provides strong evidence that alignments restricted to the primate lineage are sufficiently powerful to determine conserved secondary structural motifs in long noncoding RNAs. Given the current scarcity and challenges of obtaining experimental secondary structures of lncRNAs, we offer a promising method for identifying conserved structural motifs *in silico* through phylogenetic analyses of primate species.

## CONCLUDING REMARKS

There has been limited ability to resolve information regarding conservation of primary sequence or secondary structure of most long noncoding RNAs (lncRNAs) due to their rapid evolutionary mutation rates. We posit that a central reason why lncRNAs are considered to lack meaningful conservation of primary sequence and secondary structure was largely because of an inability to analyze a large sample size of genomes which were *similar enough*, within an appropriately narrow evolutionary span, to capture the fast rate of divergence among lncRNAs. Our approach leverages publicly available sequences of primates to address those challenges. We selected 190 unannotated primate genomes and developed a method to demonstrate the interpretability of this genomic data and publish a generalizable method. While our group studies noncoding RNA, this set of primate genomes can offer a promising approach to investigate conservation and covariance among any genomic element which evolves rapidly, including protein-coding genes or intronic regions of the genome. We ultimately demonstrate that our method can indeed resolve primary sequence conservation, the rates and spread of evolutionary divergence in the primary sequence of noncoding RNAs, and identify secondary structures in a lncRNA with significant covariance, providing proof-of-concept for the utility and applications of such a pipeline.

There are important questions which remain outstanding. Because these sequenced primate genomes have yet to be annotated or scaffolded, it remains unclear whether the observed drops in conservation corresponding to increasing divergence times are confounded by potentially poor alignment quality and a low sample size of species in the 69 MYA divergence group. It is additionally important to note the computational demand of running 190 query-to-target alignments in multi-gigabyte sized genomes. As such, there can be significant challenges with memory and computational load when aligning query RNA above 1.5-2.0 kb, as large sequences tend to require more processing power and memory, even in a high-performance computing node. Such computational strain does pose a challenge when attempting to conduct genomic analysis on long query sequences; however, this impediment can likely be mitigated by conducting multiple alignments of shorter query sequences, such as single exons, if necessary. Another limitation is that short query sequences, such as under 100 nts, may suffer from false-positive alignment results, where Exonerate may stochastically identify homologous sequences in off-target genomic loci. This limitation can be mitigated by conducting alignments with multiple concatenated exons to improve alignment quality and decrease misidentification. We ultimately hope that through an analysis of the primate lineage, novel structure-function relationships of rapidly evolving genomic machinery might yet be discovered.

## MATERIALS AND METHODS

### Genome retrieval and alignment

We developed a computational pipeline to identify homologous RNA sequences across primate genomes and generate high-coverage multiple sequence alignments (MSAs) for downstream evolutionary and structural analyses. The pipeline was designed to mine raw sequencing data from unannotated primate genome assemblies downloaded from the European Nucleotide Archive (ENA; accession PRJEB67744) (Kuderna et al. 2023). In its current implementation, the workflow was applied to 190 available primate genomes.

Genome retrieval was automated using custom Python and Bash scripts. All primate genome assemblies were downloaded from ENA and organized into a local directory structure for subsequent processing. Because many assemblies were unannotated and incompletely scaffolded, often consisting of contigs, each genome was first indexed using SAMtools (v1.23) to generate binary reference index files suitable for rapid sequence alignment (Li et al. 2009; Danecek et al. 2021). However, this indexing step was optional and not strictly necessary for downstream analyses. All scripts and instructions necessary to run the complete pipeline have been deposited at the following GitHub repository: https://github.com/pylelab/NcRNA_Evolution_in_Primates.

### Identification of putative lncRNA homologs

To identify putative homologous sequences for each query lncRNA, we performed nucleotide query-to-target pairwise alignments against each primate genome using Exonerate est2genome (v2.4.0) (Slater and Birney 2005). We used the following query sequences for our analysis: SChLAP1 isoform 3 (NCBI reference NR_104321.1; 1.1kb); P53 (NCBI reference NR_176326.1; 142 nt); Ferritin (NCBI reference NM_002032.3; 209 nt). This step generated multiple candidate alignments per genome, reflecting the fragmented structure of many assemblies. To resolve these candidates, we implemented a custom Biopython parsing script to evaluate Exonerate hits. For each genome, our script selected the highest-confidence alignment on the basis of Exonerate’s “vulgar” annotations which indicate alignment quality and coverage. Selected genomic DNA sequences were then transcribed to their corresponding RNA sequences for downstream comparative analyses.

### Multiple sequence alignment generation

The recovered RNA homologs were collated across species and aligned using MAFFT (v7.526), with the human lncRNA sequence serving as the reference anchor (Katoh et al. 2019). This produced species-level MSAs for each lncRNA or lncRNA region of interest. These alignments were used as the basis for downstream analyses of pairwise conservation, effective identity, and covariation.

### Pipeline implementation

The pipeline executes autonomously from genome retrieval through species-to-target pairwise alignment, and homolog identification. Sequence extraction, multiple sequence alignment, and downstream analyses are required to be performed by the user, though we have written custom scripts to enable these processes. By integrating ENA genome mining with established command-line bioinformatics tools and custom scripts, this framework enables systematic comparative analysis of lncRNA sequence and structural conservation across primate evolution.

### Alignment clustering for redundancy reduction

During pipeline development, we found that closely related primate species frequently yielded highly similar or identical sequences, which reduced the phylogenetic informativeness of the alignments for covariation analysis. To mitigate this, we applied CD-HIT-EST (v4.8.1) to cluster and remove near-duplicate sequences above a 95% similarity threshold prior to structural inference (word size of n=8, T=2, M=16,000) (Li and Godzik 2006). After clustering using CD-HIT-EST, the RNA sequences were then realigned using MAFFT. This redundancy reduction step improved the ability to detect compensatory substitutions indicative of conserved RNA secondary structure.

### Per-nucleotide and per-species pairwise analysis of conservation and coverage

Conservation scores were determined using Infernal (v1.1.5) in the Easel-Alistat package, and pairwise conservation of each species alignment against the human reference sequence was calculated using NCBI MSA Viewer (v1.26.0) to determine percent coverage and percent identity (Eddy 2023). Effective identity was calculated by multiplying the percent coverage by percent identity to adequately reflect the accuracy and quality of each alignment. Per-nucleotide conservation and percent non-gapped values were smoothed with a 21 nt median sliding window (step size of 1 nt).

### Mapping evolutionary divergence

The open-source database TimeTree (https://timetree.org) was used to obtain the divergence time of each species relative to *Homo sapiens* (Kumar et al. 2022). This allowed for the mapping of species conservation against divergence time to assess the rate at which RNAs diverge across the 190 primate species. Statistical significance between divergence bins was calculated by unpaired, two-tailed Student’s t-test, where **** = p ≤ 0.0001, *** = p ≤ 0.001, ** = p ≤ 0.01, and * = p ≤ 0.05.

### Covariation analysis

Covariation was evaluated using R-scape (v0.2.1) with the RAFSp command, which was applied to the filtered MSAs at a 95% identity threshold to determine compensatory substitutions which conserve base-pairing (Rivas et al. 2017; Tavares et al. 2019). To conduct analysis in R-scape, MSAs were converted from Clustal formatting (.aln) to Stockholm format (.sto) using ConvertAlign (Center for Integrative Bioinformatics VU, https://www.vubioinformatics.com/tools/#msa). Together, these analyses enabled the detection of putatively functional secondary structural motifs within primate lncRNAs.

### Preparing SChLAP1 secondary structure for covariation analysis

We obtained the *in cellulo* SHAPE-Map secondary structure reactivities of SChLAP1 isoform 1 replicate 1 (Falese et al. 2025). We predicted the secondary structure of isoform 1 by passing the SHAPE-Map reactivities into SuperFold under default parameters (Siegfried et al. 2014; Smola et al. 2015). After converting the .ct to a dot-bracket structure file, we aligned the primary sequence of SChLAP1 isoform 1 to SChLAP1 isoform 3. We identified that SChLAP1 isoform 1 has an insertion from nucleotides 437-772. Thus, we removed positions 437-772 from the dot bracket file to align the structure file to isoform 3. We also identified 55 nucleotide positions which were base pairing into the removed region and converted those nucleotide positions into dots (single-stranded) in the dot-bracket structure to ensure that the number of open-brackets was equal to the number of closed-brackets in the structure file. This provided a structure file for SChLAP1 isoform 3. Because all the MSAs contained gaps and insertions, and because the secondary structure file entered into R-scape must be equal in length to the number of alignment columns, it was then necessary to add gaps into the secondary structure file as dots in the dot-bracket structure, which we performed for each alignment with a custom Biopython script. After obtaining the proper secondary structure for each alignment, we added it to the Stockholm file for input into R-scape.

## DATA DEPOSITION

Processed conservation and covariation data are available as Supplemental Data files. All scripts and instructions necessary to run the complete pipeline have been deposited at the following GitHub repository: https://github.com/pylelab/NcRNA_Evolution_in_Primates.

## Supporting information

Supplemental_Data_1_Conservation_and_Coverage

Supplemental_Data_2_Pairwise_Divergence

Supplemental_Data_3_Covariation

## ACKNOWLEDGEMENTS

Thank you to the Yale Center for Research Computing for their guidance and use of the HPC clusters. A.M.P. is an investigator of the Howard Hughes Medical Institute. This work was also supported by the NIH (1R01HG011868-01 to A.M.P.) and the NSF GRFP (DGE-2139841 to Z.R.P.).

## COMPETING INTEREST STATEMENT

None to declare.

## Notes

https://github.com/pylelab/NcRNA_Evolution_in_Primates

### Competing Interest Statement

The authors have declared no competing interest.

